# A seleno-hormetine protects bone marrow hematopoietic cells against ionizing radiation-induced toxicity

**DOI:** 10.1101/432286

**Authors:** Bartolini Desirée, Wang Yanzhong, Zhang Jie, Giustarini Daniela, Rossi Ranieri, Gavin Y. Wang, Torquato Pierangelo, Danyelle M. Townsend, Kenneth D. Tew, Galli Francesco

## Abstract

2,2’-diselenyldibenzoic acid (DSBA) is a mild thiol peroxidase agent presently in preclinical development. This study reports that the drug has novel seleno-hormetic properties in both murine bone marrow and human liver cells. According with previous in vitro findings, mechanistic aspects of such properties were confirmed to include the activation of Nrf2 transcription factor and an increased expression of downstream stress response genes in the liver and in hematopoietic stem and progenitor cells of the myeloid lineage. These genes include glutathione S-transferase that is reported to represent a major player in the metabolism and pharmacological function of seleno-organic compounds. As a practical application, DSBA administration prevented bone marrow toxicities following acute exposure to sub-lethal doses of ionizing radiation in C57 BL/6 mice.

In conclusion, this study demonstrates for the first time the pharmacological properties of DSBA *in vivo*. The findings suggest applications for this selenohormetine in radioprotection and prevention protocols.

## 1. Introduction

Inorganic and organic forms of selenium have extensively been investigated as pharmacological agents with applications in either cancer chemoprevention (cytoprotective effects) or therapy of drug-resistant tumors (recently reviewed in [1]). These compounds act as thiol peroxidases (TP) and agonists of drug metabolism genes associated with the detoxification of cellular electrophiles [2]. Recently, we demonstrated that structural modifications of the diphenyldiselenide [(PhSe)_2_] scaffold, can lead to mitigation of the redox cycling activities of the Se–Se functional group of this potent Se-organic thiol peroxidase (SeTP) [3], thus lessening its cytotoxicity [4]. 2,2’-diselenyldibenzoic acid (DSBA) is the resultant diselenide generated by this strategy that possesses *in vitro* pharmacological properities and insignificant toxicity. However, its TP activity is sufficient to stimulate an efficient adaptive redox stress response that increases protection against H_2_O_2_-induced injury in both murine embryonic fibroblasts and human hepatocytes [4]. The hormetic effect of DSBA involves the activation of the transcription factor NF-E2-Related Factor 2 (Nrf2) and is influenced by the expression of the isoform P of the enzyme glutathione S-transferase (GSTP). GSTs are among the most abundant Cys-containing cellular proteins of the liver and were the first identified to react with Se-organic compounds thus promoting their hepatic metabolism [5,6]. In this context, recent studies by some of us demonstrated that the GSTP isoform is critical for detoxification and maintenance of redox homeostasis in cells treated with SeTP [1]. The importance of this protein in regulating cellular signaling events and in initiating response to oxidative stress has been reported in some detail and implies that it may act in concert with Nrf2 to regulate a variety of cellular pathways [7].

In the present study, *in vitro* results are extended into an animal model of oxidant stress by acute exposure to ionizing radiations (IR), to examine whether DSBA had sufficient hormetic activity to prevent damage to hematopoietic stem and progenitor cells from bone marrow (BM), a major biological consequence of radiation exposure in this model [8].

## 2. Materials and Methods

### 2.1 Seleno-Compounds

2,2’-diselenyldibenzoicacid (DSBA) was synthesized as reported in [4]. Purity > 98.5%.

Ebselen (E3520) and diphenyl-diselenide[PhSe)_2_] (180629; purity 98%) were purchased from Sigma-Aldrich and all compounds were dissolved in DMSO.

### 2.2 In vitro studies in human liver cell lines

HepG2 human hepatocarcinoma cells were maintained in MEM medium (Gibco, Life Technology) supplemented with 10% fetal bovine serum (Gibco, Life Technology) in the presence of 100 U/ml penicillin and 100 mg/ml streptomycin (Sigma-Aldrich, USA). HepaRG human progenitor hepatic cells (Thermo Fisher Scientific) were maintained according to the manufacturer’s recommendations. Briefly, the cells were grown in William’s E medium (Thermo Fisher Scientific) supplemented with Glutamax (Gibco), 5 ug/mL human insulin (Sigma-Aldrich) and 50 µM Hydrocortisone hemisuccinate (Sigma-Aldrich) for 14 days. All cells were kept at 37°C in a humidified 5% CO_2_ cell culture incubator and were passaged using trypsin-EDTA (Euroclone).

### 2.3 Cellular thiols and glutathionylation

Cellular thiols were assessed by HPLC analysis with fluorescence detection after derivatization with monobromobimane (mBrB, Calbiochem). For disulfide analysis, aliquots of samples were derivatized with N-ethylmaleimide (Sigma-Aldrich) to mask reduced thiols and then dithiothreitol (DTT, Sigma-Aldrich) was used to reduce disulfide bridges, according to Rossi et al. [9].

The Cayman’s Glutathionylated protein detection kit (Cayman Chrmical, Item No.10010721) was used to assess PSSG in MEFs. The method allows a direct measurement of *S*-glutathionylated proteins in whole (permeabilized) cells by flow cytometry (Attune NxT Acustic Focusing Cytometer, Thermo Fisher Scientific).

### 2.4 In vivo and ex vivo studies

Male C57 BL/6 mice purchased from the Jackson Laboratories (Bar Harbor, ME) and were used for *in vivo* experiments. The animals were housed five per cage in the Hollings Cancer Center AAALAC-certified animal facilities at the Medical University of South Carolina (MUSC). Animals received food and water ad libitum. All mice were used at approximately 8 –12 weeks of age. The Institutional Animal Care and Use Committee of MUSC approved all experimental procedures used in this study. DSBA was dissolved in DMSO and then diluted with 30% PEG2000/PBS. Mice were administered with a single dose of the diluted DSBA solution at 10 mg/Kg and 50 mg/Kg via intraperitoneal injection. Control animals were treated with the vehicle. The groups of mice included 3 animals each. Mice were scarified 24 hrs after the treatment to collect blood, bone marrow (BM) and liver samples.

### 2.5 Total-Body Irradiation (TBI) and DSBA treatment

To investigate the protection against IR injury, the number of BM HSPCs was evaluated in animals that received a dose of 50 mg/kg DSBA 4 h before TBI exposure. Mice were exposed to 3 Gy of irradiation using a J. L. Shepherd Model 143 ^137^Cs gamma irradiator at a dose rate of 2.0 Gy/min as described previously [10]. Twenty-four hours after TBI, mice were euthanized by CO_2_ suffocation followed by cervical dislocation, and the femora and tibiae were immediately harvested from the mice for the isolation of bone marrow mononuclear cells as described below.

### 2.6 Isolation of BM Mononuclear Cells (BM-MNCs)

The femora and tibiae were harvested from the mice immediately after they were euthanized with CO_2_. Bone marrow cells were flushed from the bones into Hank’s buffered saline solution (HBSS) containing 2% FCS using a 21-gauge needle and syringe. Cells from at least three mice were pooled and centrifuged through Histopaque 1083 (Sigma, St. Louis, MO) to isolate bone marrow BM-MNCs as described previously [10].

### 2.7. Flow Cytometric analysis of Hematopoietic Cells

Flow cytometry was used to analyze Hematopoietic Stem Cells (HSCs) and Progenitor cells (HPCs) as previously described [11]. Briefly, BM-MNCs were incubated with PE-conjugated antibodies against CD3e, CD45R/B220, Gr-1, Mac-1 and Ter-119 to stain the lineage-positive cells. The cells were washed with PBS and incubated with anti-CD16/CD32 antibody to block Fc receptors. Finally, the cells were stained with PE-Cy7 conjugated anti-Sca-1 and APC-H7 conjugated anti-c-kit antibodies and analyzed using a BD LSRFortessa^™^ X-20 flow cytometer (Becton Dickinson, San Jose, CA). The data were analyzed using FlowJo software. Cells stained negative for lineage markers and c-kit but positive for Sca1 were considered as HPCs (lineage^-^/Scal1^-^/c-kit^+^cells, or LSK^-^ cells) and those negative for lineage markers but positive for Sca1 and c-kit as HSCs (lineage-/Sca1^+^/c-kit^+^ cells, or LSK cells).

### 2.8 Flow cytometric analysis of ROS

ROS levels were measured in HSCs and HPCs as an *in vivo* indicator of DSBA toxicity by Flow Cytometry using the probe DCFH-DA. Intracellular ROS were measured by flow cytometric analysis as previously reported [8]. Briefly, Lin^-^ HSPCs were loaded with 5 mM of DCF-DA and incubated at 37 °C for 30 min. The levels of ROS in HSPCs were analyzed by measuring the mean fluorescence intensity of DCF-DA using a BDLSRFortessa^™^X-20 cell analyzer (Becton Dickinson, San Jose, CA) and FACSDiva^™^ software. Data analysis was performed using FlowJo software (Tree Star, Ashland, OR).

### 2.9 Colony-forming unit assay

Colony-forming unit (CFU) assays were performed by culturing the isolated BM-MNCs in MethoCult GF M3434 methylcellulose medium (Stem Cell Technologies) as described previously [12]. Colonies of colony-forming unit-granulocyte macrophage (CFU-GM) and burst-forming unit-erythroid (BFU-E) were scored on day 7, while colonies of CFU-granulocyte,–erythrocyte,–monocyte, and -megakaryocyte (CFU-GEMM) were enumerated on day 12 after incubation.

### 2.10 Nuclear and cytosolic protein extraction

Cellular extracts obtained after 4 h of treatment with DSBA (from 5 to 20 µM) were used for these experiments and cellular proteins were probed before or after fractionation of cytosolic and nuclear proteins carried out utilizing a Thermo Scientific NE-PER Nuclear and Cytoplasmic Extraction Kit (Cat# 78833, Thermo Fisher).

### 2.11 Western blotting analysis

Protein samples were extracted using cell lysis buffer (Cell Signaling) supplemented with a cocktail of proteinase inhibitors (Sigma) and protein concentrations were determined using the Bio-Rad Dc protein assay kit (Bio-Rad Laboratories). Western blots was performed as described in [13]. Briefly, 50 µg of protein samples were resolved on 10% Mini-Protean TGX gels (Bio-Rad) and transferred onto 0.2 mM PVDF membrane (Millipore). Blots were blocked with 5% nonfat milk for 1–2 h at room temperature, then probed with primary antibodies, and incubated at 4 °C overnight. Primary antibodies used were: anti-GSTP, anti-Nrf2 (#12721) and anti-aldehyde dehydrogenase-1 (ALDH1) (#12035) from Cell Signaling; heme-oxygenase 1 (HO-1) (SC-390991), anti-Nrf2 (SC-772) and Tubulin (SC-23948) from Santa Cruz Biotechnology. After extensive washing with TBST, blots were incubated with appropriate HRP-conjugated secondary antibody for 1.5 h at room temperature.

Protein bands were detected using an ECL Plus Western Blot Detection System (GE Healthcare Life Science).

### 2.12 GST activity

The specific activity of the GST in bone marrow and plasma samples were measured as previously described in [14] using 5 mM GSH (Sigma-Aldrich, St. Louis, MO) and 0.5 mM CDNB (Merck, Darmstadt, Germany) as second substrate in 0.1M potassium phosphate buffer pH 6.5 at room temperature with the Benchmark plus microplate spectrophotometer (BioRad, Hercules, CA) by following the change in absorbance at 340 nm. The molar extinction coefficient used for CDNB conjugation was 9.6 mM^-1^cm^-1^. Enzymatic activities were calculated after correction for the non-enzymatic reaction.

### 2.13 Immunohistochemical analysis (IHC)

Hepatic Nrf2 was measured by immunohistochemistry (IHC) as previously described in [15]. Briefly, mouse liver tissues were fixed with formalin and embedded in paraffin. Tissue sections (5 μm thick) were prepared. Endogenous peroxidase activity was blocked by incubation with 3 % hydrogen peroxide for 30 min and followed by heating in 1mM EDTA for antigen retrieval. The sections were then blocked with 5 % normal goat serum in 0.1 % Triton X-100/PBS for 1 h and incubated overnight at 4 degree with rabbit anti-human Nrf2 antibody (1:200, Santa Cruz). After wash with PBS, slides were incubated with ABC reagent (Vector) for 30 min. Immunostaining was visualized by DAB and the slides were counterstained using hematoxylin.

### 2.14 Statistics

Data (as means+/−SD) were assessed for distribution and differences between variables were assessed for statistical significance using parametric or non-parametric tests when appropriate.

## 3 Results

### 3.1 In vitro effects of DSBA on liver cell ROS and thiols

To further characterize the liver cell metabolism and redox function of DSBA, we comparatively assessed the effect of this and other Se-organic drugs on ROS generation and thiol levels of HepaRG hepatic stem cells and HepG2 hepatocarcinoma cells. DSBA was less effective than Ebselen or the diselenide precursor (PhSe)_2_ in stimulating the generation of cellular ROS that was generally higher in HepaRG than in HepG2 cells (Figure 1). Accordingly, a slight increase in the generation of cellular ROS was only observed when HepaRG cells were treated with 50 µM DSBA that is the highest concentration of the compound tested in this study (Figure 1). In both the cell lines, treatments with DSBA and the other SeTP at a final concentration of 10 µM did not cause significant reductions of cell viability (not shown).

**Figure 1.**
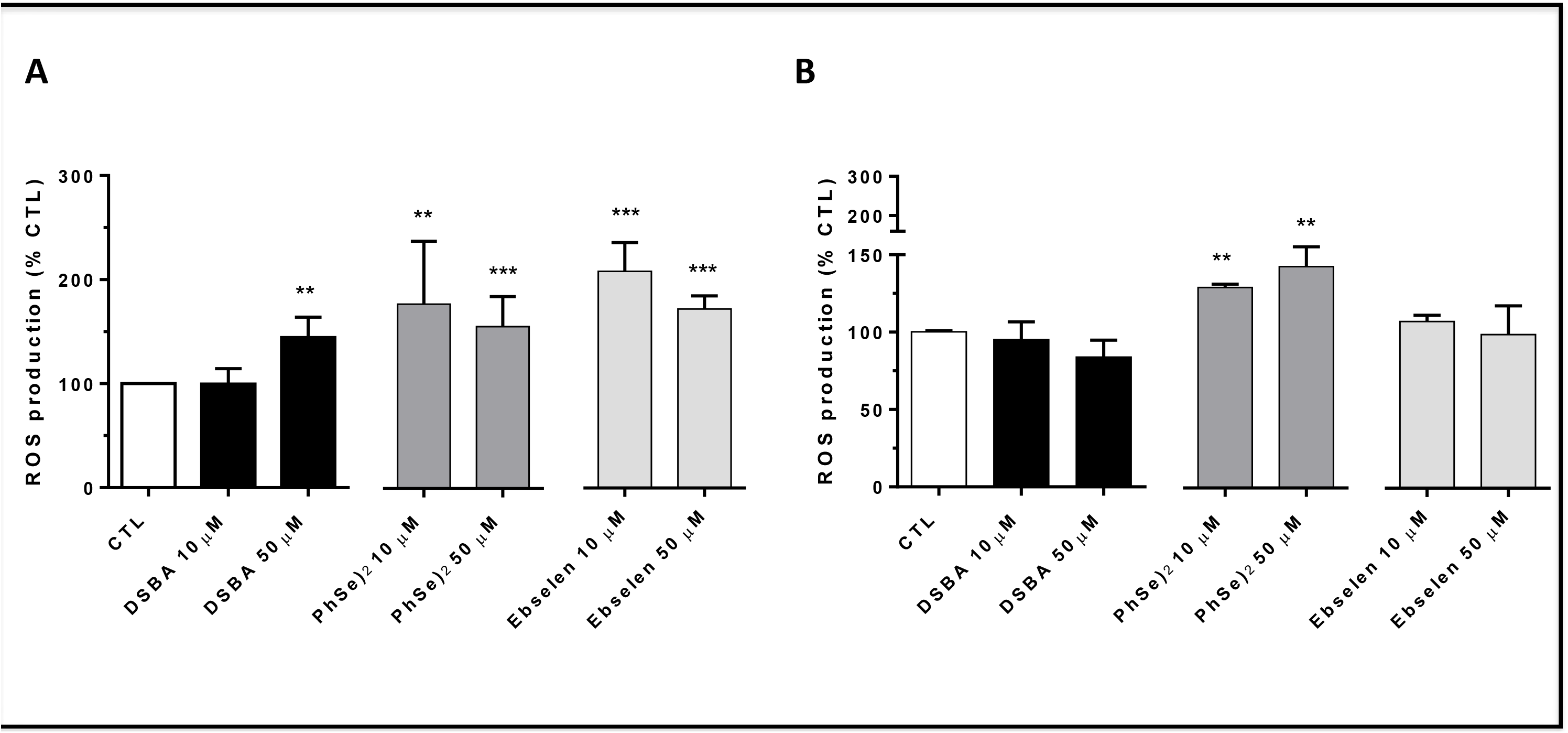
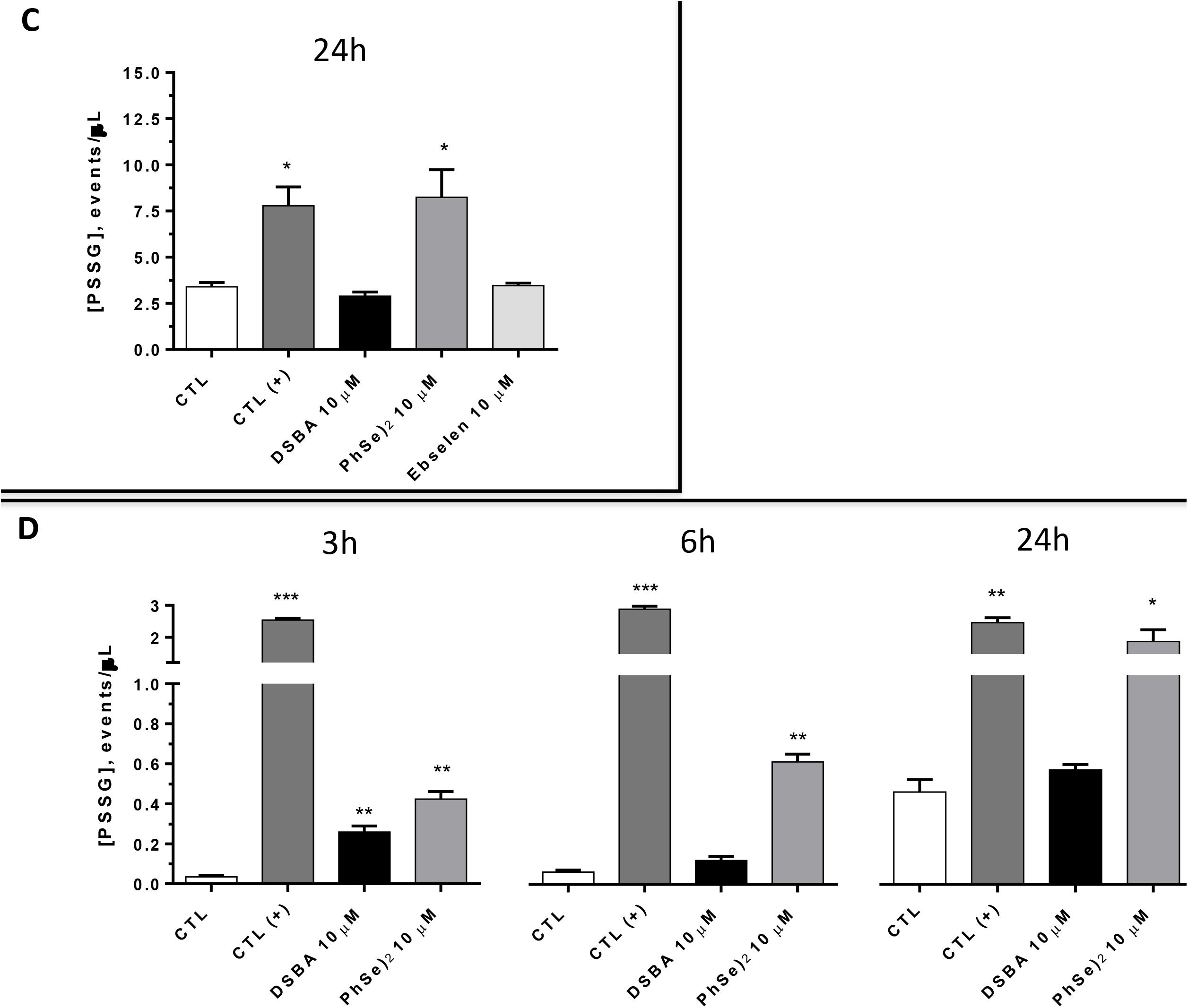
Reactive oxygen species (A and B) and protein S-glutathionylation (C and D) in HepaRG and HepG2 cells. HepaRG or HepG2 cells were alternatively treated for 24 hours with 10 μM DSBA, PhSe)_2_ or Ebselen then ROS were measured the DCF method. PSSG were assessed in permeabilized cells by FACS-Scan as described in the text.

Lower fluxes of ROS in HepG2 hepatocarcinoma cells compared to HepaRG cells might be explained by the higher cellular levels of GSH (Table1) and average GSH/GSSG ratio (256 and 170, respectively; p < 0.05). This more reduced environment of HepG2 cells was sustained by the lower capability of GSH secretion in the extracellular medium (Table1) and protein S-glutathionylation (Figure 1). In these human hepatocarcinoma cells, DSBA did not change the intra-and extra-cellular levels of GSH, as well as the cellular levels of GSSG (Table1), but did significantly increase PSSG levels (Figure 1). Conversely, the more potent TP compound Ebselen [3], stimulated the GSH metabolism of HepG2 cells, increasing its cellular levels and secretion, and its oxidation to form PSSG (Figure 1) and GSSG (Table 1) at the cellular level.

**Table 1.** Cellular and extracellular levels of glutathione in tumoral HepG2 and non-tumoral HepaRG human liver cell lines treated for 24 hours with DSBA.

|  |  |  |  |  |
| --- | --- | --- | --- | --- |
|  | Intracellular GSH<br>(nmol/10<sup>6</sup> cells) | Extracellular GSH<br>(nmol/10<sup>6</sup> cells) | Intracellular<br>GSSG<br>(nmol/10<sup>6</sup> cells) | GST activity<br>(U/mg of proteins) |
| **HepG2 cells** |  |  |  |  |
| CTL | 33.3 ± 5.37 | 2.25 ± 1.41 | 0.13 ± 0.03 | 53.4 ± 6.64 |
| DSBA 10 µM | 31.5 ± 5.12 | 3.32 ± 1.50 | 0.15 ± 0.05 | 57.2 ± 0.65 |
| DSBA 50 µM | 27.2 ± 6.2 | 2.48 ± 1.2 | 0.12 ± 0.04 | 41.0 ± 10.03* |
| Ebselen 10 µM | 45.0 ± 5.0 | 4.40 ± 1.31 ** | 0.18 ± 0.05 * | 23.2 ± 8.31** |
| Ebselen 50 µM | 50.3 ± 4.2 * | 5.06 ± 2.0 ** | 0.40 ± 0.06 ** | 43.1 ± 2.21* |
| **HepaRG cells** |  |  |  |  |
| CTL | 18.7 ± 2.12 | 10.25 ± 3.70 | 0.11 ± 0.01 | 18.2 ± 0.63 |
| DSBA 10 µM | 25.60 ± 4.3 * | 13.85 ± 5.2 | 0.13 ± 0.03 * | 6.8 ± 0.19 * |
| Ebselen 10 µM | 18.80 ± 5.0 | 16.54 ± 4.31 * | 0.18 ± 0.07 ** | 16.3 ± 0.07 |
| t-test: CTL (vehicle = DMSO) vs treatment *p<0.05; **p<0.01.<br>Data in parentheses are the percentage of activity calculated on the CTL. |  |  |  |  |

In HepaRG cells, DSBA, but not Ebselen, significantly increased intracellular GSH (Table1). This resulted in changes of the GSH/GSSG ratio (from 256 to 200), while a more marked decrease was found in Ebselen treated cells that showed an average value of 100. Under these conditions neither compound stimulated a significant increase in PSSG (Figure 1).

DSBA inhibited GST activity in HepaRG cells, but not in HepG2 cells (Table1); the latter cell type, on the contrary, showed GST activity inhibition when treated with Ebselen.

### 3.2 DSBA activates liver tissue Nrf2 in vivo

Acute exposure to DSBA at the doses used in this study did not cause overt toxicity as demonstrated by objective examination of animal behavior and clinical cues, liver histology (Figure 2), and blood and BM cellular composition and morphology (not shown). IHC analysis revealed that Nrf2 expression levels were markedly increased in mouse liver tissue in a dose-dependent fashion after DSBA injection, showing that DSBA treatment may activate Nrf2 in animal tissues in vivo (Figure 2).

**Figure 2.**
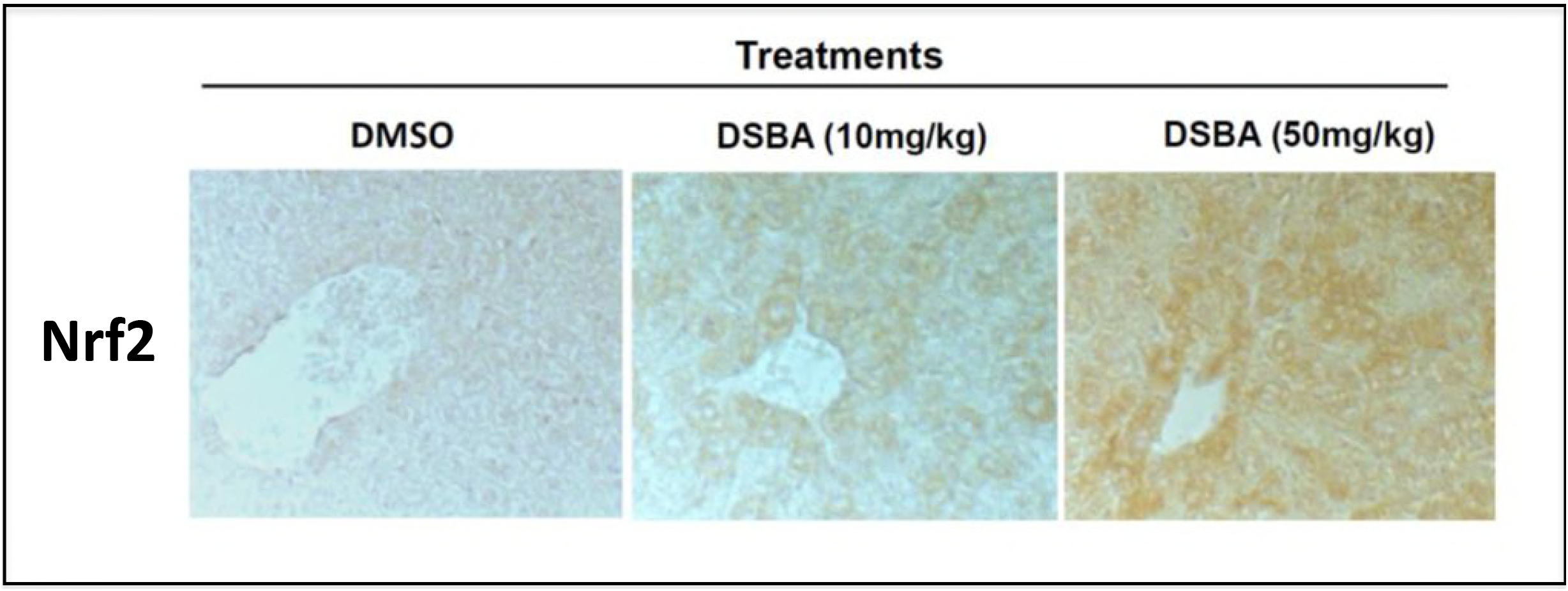
DSBA activates Nrf2 in liver tissues in vivo. IHC was employed to assess Nrf2 expression in liver tissues of C57 BL/6 mice at 24 h after drug treatment. Magnification 400x. Vehicle control = DMSO.

### 3.3 DSBA modulates the redox signaling of HSPCs in vivo

Our previous studies have shown that DSBA regulates redox status in various cells *in vitro* (6). However, it remains to be determined if DSBA affects redox balance in tissues and cells *in vivo*. As such, we investigated the impact of DSBA on ROS levels in HSPCs of C57 mice. Flow cytometry data showed that DSBA stimulates ROS generation in both HSCs and HPCs (Figure 3); reaching peak effect at a dose of 10 mg/kg and decreasing at 50 mg/kg.

**Figure 3.**
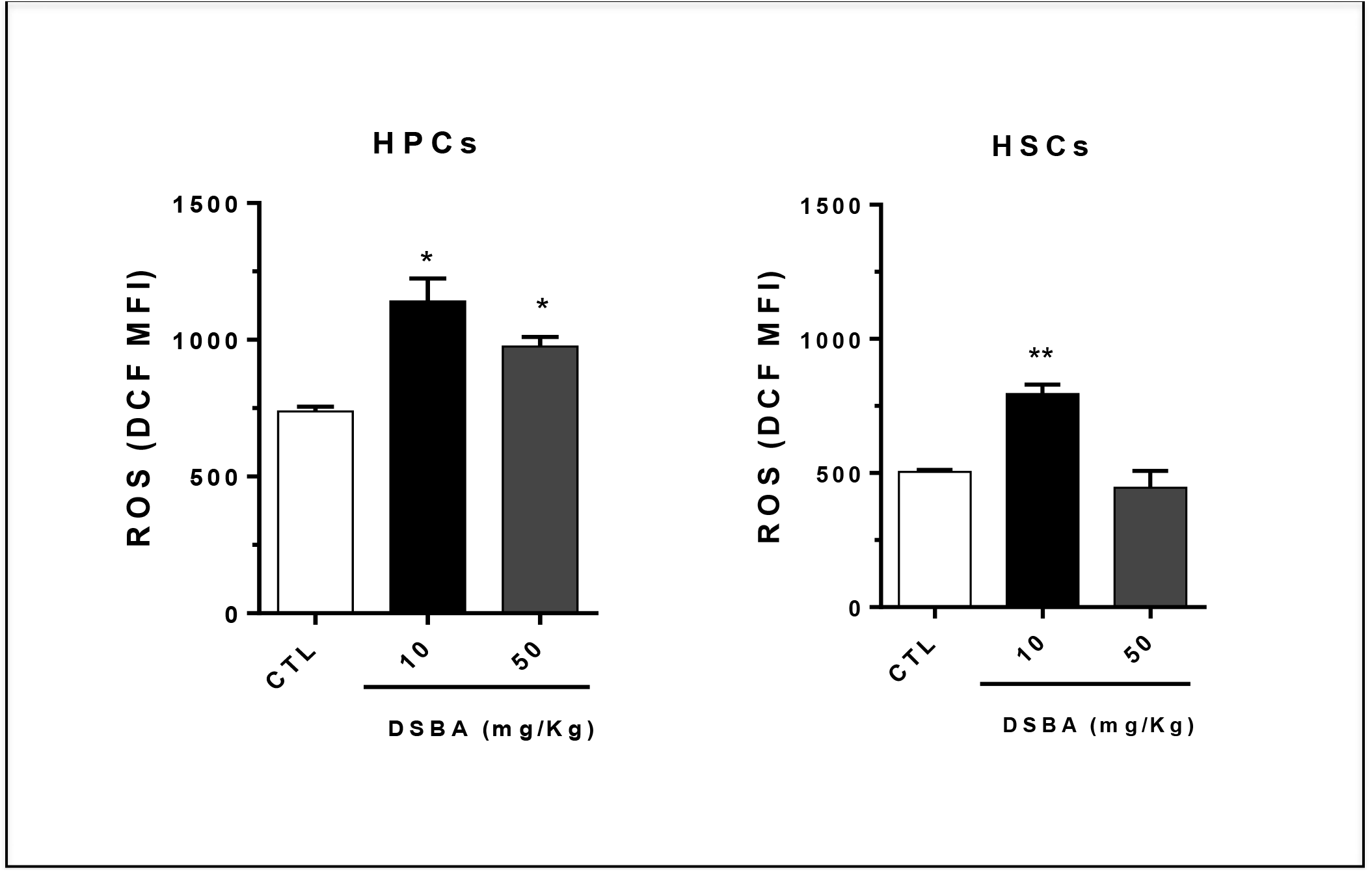
Levels of reactive oxygen species (ROS) in HPCs and HSCs isolated from DSBA-treated C57 BL/6 mice. BM-MNCs were collected at 24 h after DSBA treatment and subjected to immune-phenotype assays. DCF-DA staining and flow cytometric analysis were performed to measure ROS levels in HSCs and HPCs as described previously (8). t-test: * p < 0.05, ** p < 0.01.

Since GST plays a significant role in maintaining redox balance, we examined whether DSBA-mediated increases in ROS were due to inhibition of GST activity. Our data showed that there was no significant change in plasma GST activity after DSBA treatment (Figure 4A), but such activity significantly increased in BM-MNCs obtained from animals treated with 50 mg/kg DSBA (Figure 4B). These results imply that DSBA-induced increases in ROS are not likely the consequence of GST inhibition. In contrast, DSBA-mediated ROS production may trigger a cellular adaptive response and consequently a slight increase of GST activity in HSPCs.

**Figure 4.**
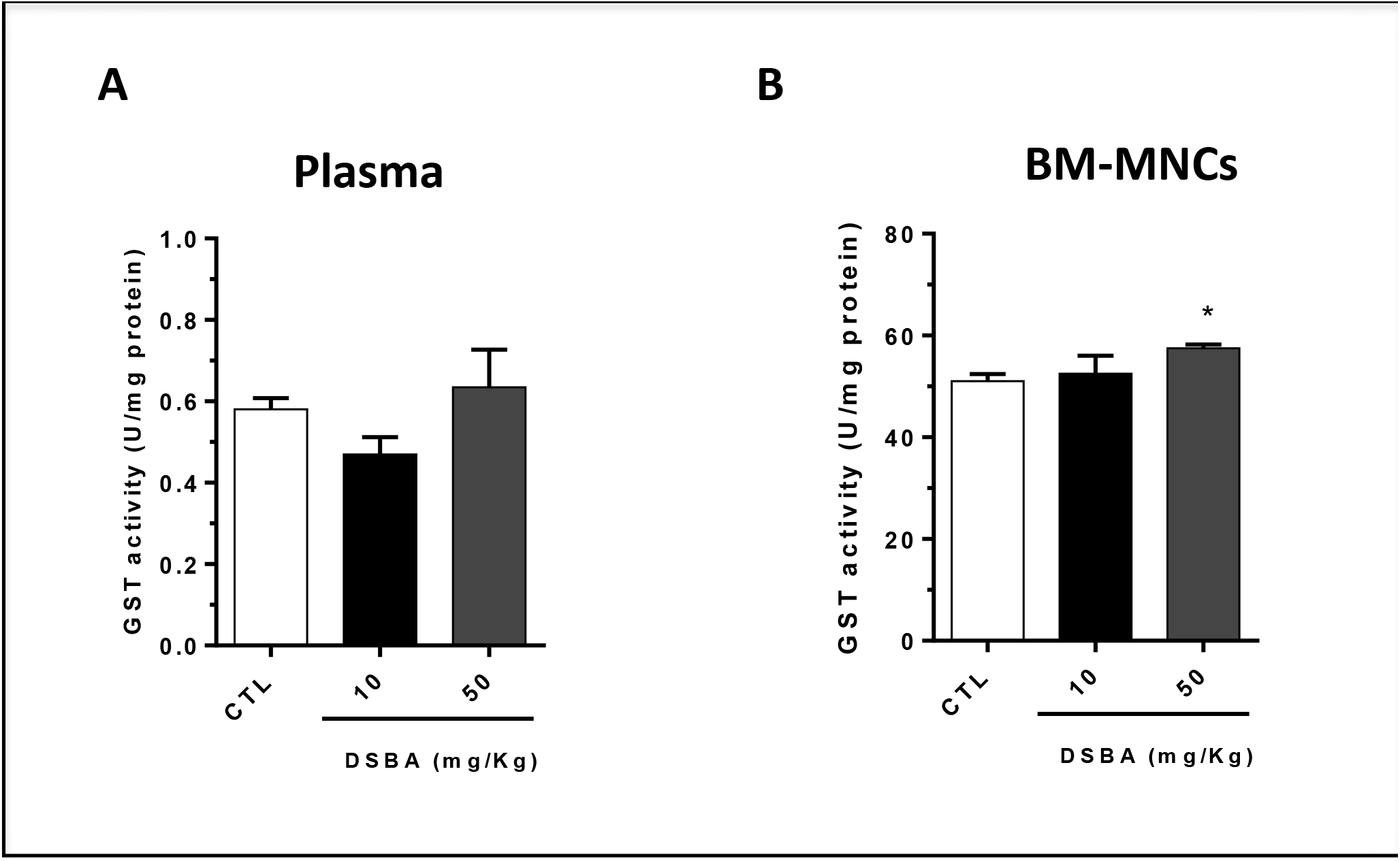
GST activity in C57 BL/6 mice plasma (A) and BM-MNCs (B) after DSBA treatment. GST activity in plasma and BM-MNCs was measured at 24 h after DSBA treatment using approch as described in the methods section. t-test: *p<0.05.

### 3.4 Hematopoietic radioprotection by DSBA correlates with Nrf2 activation in BM-MNCs

Nrf2 is a master modulator of cellular antioxidant response transcriptionally regulating expression of a variety of cytoprotective genes. To elucidate the mechanisms by which DSBA protects HSPCs against radiation injury, we determined whether this compound impacts Nrf2 signaling in BM hematopoietic cells. Immunoblot of BM-MNC proteins further confirmed the *in vivo* effects of DSBA as an Nrf2 activator; besides Nrf2 transcription (forward-feeding, self-regulating), DSBA treatment increased expression of GSTP and the other Nrf2-dependent genes HO-1 and ALDH1 (Figure 5A); at the same time, Nrf2 protein levels slightly increased after DSBA treatment in the nuclear fraction of BM-MNCs (Figure 5B); ALDH1 and GSTP proteins were also present in the nucleus and their levels increased following DSBA (Figure 5B).

**Figure 5.**
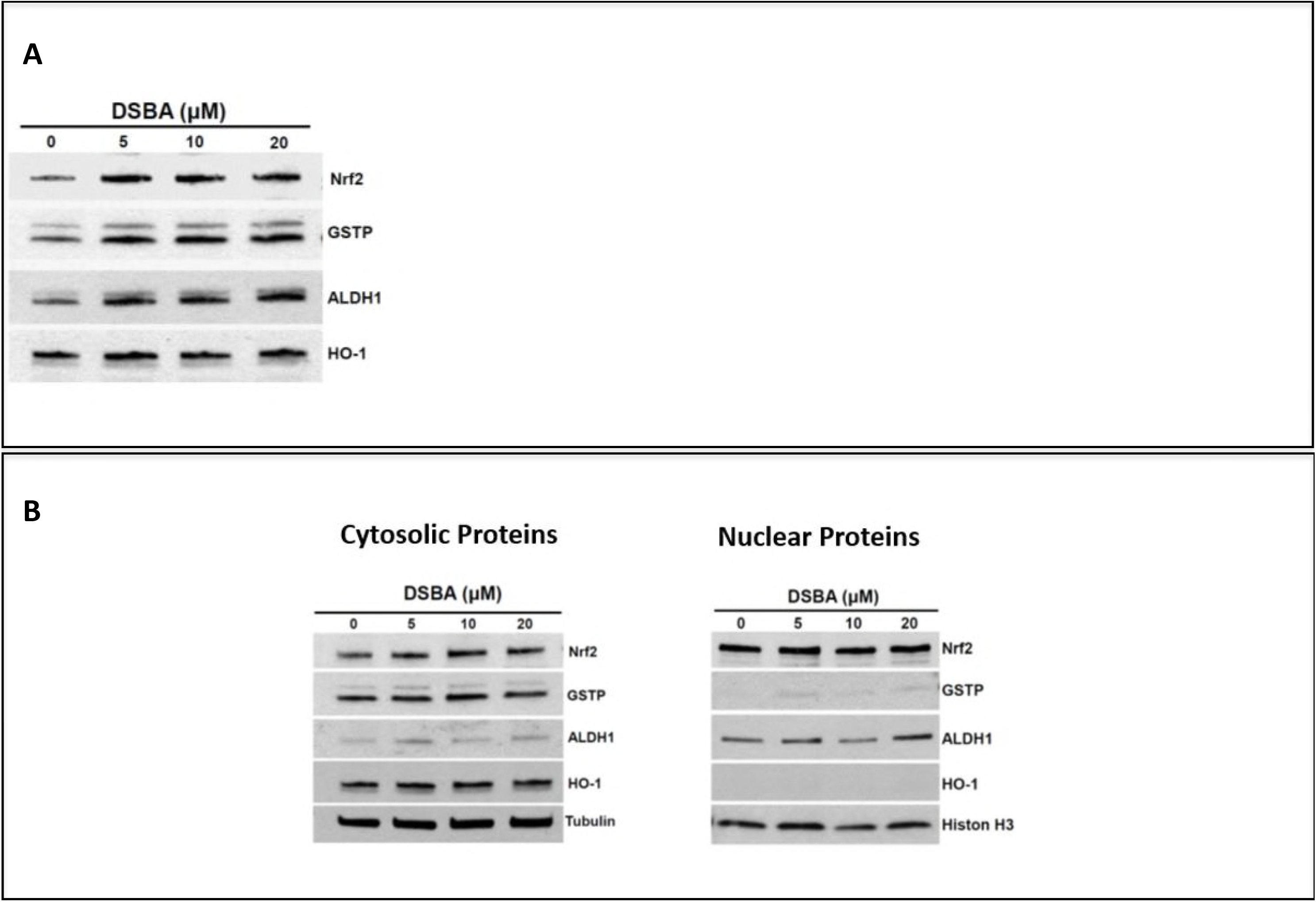
Transcriptional activation (A) and nuclear translocation (B) of Nrf2 in DSBA-treated BM-MNCs. Cells were treated for 4h with DSBA at increasing concentrations from 5 to 20 R$ and levels of Nrf2 protein and the Nrf2-dependent genes HO-1, ALDH1, and GSTP were assessed by immunoblot in cellular extracts before (A) or after fractionation of cytosolic and nuclear components (B).

### 3.5 DSBA protects HSPCs against radiation injury in vivo

The hematopoietic system is highly sensitive to radiation injury and a dose beyond 2 Gy may lead to BM suppression characterized by neutropenia, lymphocytopenia and thrombocytopenia. In light of the Fukushima nuclear accident and the increasing risk of radiation-induced BM injury, there is a critical need to develop new countermeasure agents against radiation-induced toxicity. To explore the therapeutic potential of DSBA as a new radiation protector, we treated C57 mice with this compound before exposing animals to TBI. CFU assays were employed to measure the colony-forming capacities of HSPCs. The results showed that DSBA pre-treatment prevented the IR-induced decrease of CFU-GM, BFU-E and CFU-GEMM numbers (Figure 6), indicating that DSBA does possess radioprotective properties against IR-induced injury in HSPCs.

**Figure 6.**
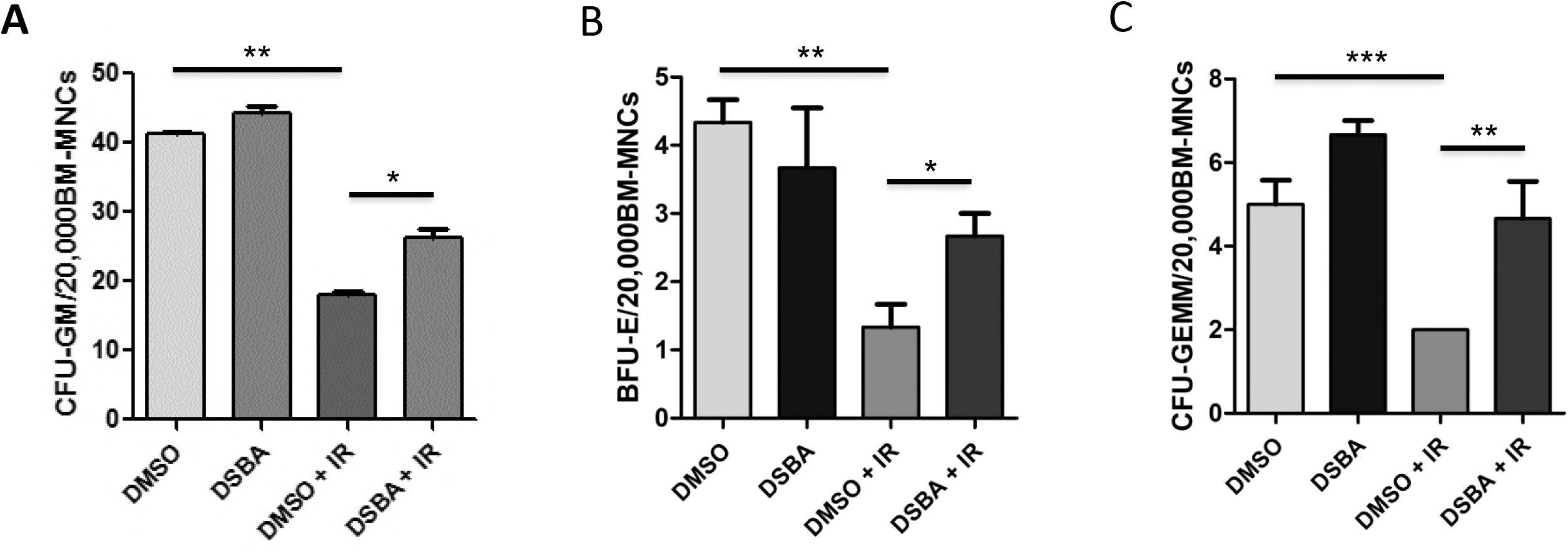
DSBA pre-treatment protects mouse BM HSPCs against ionizing radiation (IR)-induced injury in vivo. The clonogenic function of HSPCs was measured using CFU assays (12). average number of (A) CFU-GM, (B) BFU-E and (C) CFU-GEMM in 20,000 BM-MNCs.

## 4 Discussion

The findings in this study confirm the recently identified *in vitro* activity of DSBA as an Nrf2-acivating seleno-hormetine [4] *in vivo*. Nrf2 activation was demonstrated in the present study assessing liver tissue and BM HSPCs following sub-cytotoxic concentrations of DSBA in C57 BL/6 mice. The canonical Nrf2 activation model predicts that DSBA generated ROS (in both hepatocytes and BM) stimulates this transcription factor through dissolution of its interaction with Keap1, allowing migration to the nucleus promoting antioxidant and electrophile responsive elements [16]. Indeed, the lowest dose of DSBA investigated in this study caused such a response in BM cells, associated with nuclear translocation of Nrf2 protein and expression of a series of Nrf2-dependent genes. In these BM cells, GSTP was found to be the most responsive gene followed by ALDH1 and then HO-1, and GSTP and ALDH1 were also upregulated in the nucleus. In mice treated with 50 mg/kg DSBA, such gene response was associated with lowered ROS levels, implying a rapid and efficient detoxification response mediated through Nrf2 activation.

The observation that DSBA increased hepatic Nrf2 *in vivo* (Figure 2) is in agreement with previous findings obtained in human liver cells [4]. Since this transcription factor plays a role in hepatic and systemic metabolism of GSH and Cys [16], and thiol-mediated pathways are implicated in Se-compound detoxification [1], the influence of DSBA on liver cell glutathione was investigated. DSBA increased cellular GSH and ROS flux in HepaRG cells with minor effects on GSH secretion into the extracellular milieu and oxidation to GSSG, effects that are commonly associated with the exposure to Se-compounds [17] and other electrophiles [18]. The more reduced intracellular environment of HepG2 cells prevented these effects, with only slightly increased levels of protein S-glutathionylation, probably dependent upon higher levels of GSTP in this hepatocarcinoma cell line. Conversely, treatment with other compounds with much higher TP activity, i.e. Ebselen and (PhSe)_2_, markedly influenced the redox homeostasis of these liver cells. We therefore demonstrated in this study that DSBA is a relatively safe Se-organic molecule with minor effects on liver cell redox and low toxicity, even though the drug effects are sufficient to activate Nrf2 and its detoxification gene response both *in vitro* and *in vivo*.

DSBA was also confirmed to inhibition GST activity in HepaRG cells and to stimulate GSTP gene expression *in vivo*. This is not a trivial observation if we consider that GST is the first and likely the most important thiol-containing protein identified to react with SeTP, and Cys alkylation of GST protein is demonstrated promoting Se-compound sequestration and metabolism in the liver [5,6]. This alkylation reaction produces the irreversible inhibition of GST enzyme activity, a process that was originally characterized for the prototypal compound Ebselen [6] and a response that is confirmed to occur in the in vitro experiments on liver cells of this study. Alkylating agents are potent GST gene inducers and also a cause of drug resistance [1,19]. GST-overexpressing tumor cells, such as the hepatocellular carcinoma HepG2 cell line investigated herein, are in fact poorly responsive to the mild thiol peroxidase activity of DSBA while the reactive compound Ebselen provokes a significant inhibition of the enzyme activity, then leading to increased GST gene and protein expression (previously characterized in [3]). In these cells and in other cell models, the GSTP isoform was demonstrated to be particularly important in provide an efficient cellular response to either the hormetic or cytotoxic activity of SeTP [3,4]. Intriguingly, such response appears to depend on the capability of GSTP isoform to functionally and physically interact with Nrf2 protein during the gene induction process [4].

At the same time, GSTP gene expression is enmeshed in pathways that control proliferation and migration of BM myeloid cells and among these cells, the myeloid lineage is known to be highly responsive to GSTP-targeted pharmacological agents [20]. Intriguingly, *in vivo* treatment with DSBA increased GSTP expression both in the cytosol and the nucleus of BM progenitor cells. This finding is in agreement with the previously reported co-localization of GSTP and Nrf2 in both the cytosolic and nuclear compartments during drug-induced activation [4]. Therefore, the nuclear availability of GSTP together with relevant concentrations of protein thiols and GSH in the nuclear environment [21], make such co-localization potentially strategic for nuclear protection and redox-dependent regulation of transcriptional processes associated with SeTP detoxification.

A novel component of this study was to determine whether DSBA may have *in vivo* hormetic effects. A total body irradiation model was used for these experiments which is associated with oxidative stress, causative of BM stem cell damage and subsequent myelosuppression [8]. Our results conclusively demonstrated that DSBA pretreatment prevents hematopoietic stem cell damage and death in IR-exposed animals. In this regard, recent studies demonstrated that some redox-active superoxide dismutase mimics produce similar positive effects on BM cells and behaving mechanistically in the same fashion [22]. Therefore, these results may be expanded to consider whether DSBA can be used to manage radiation emergency situations or the types of hematologic toxicity observed in patients undergoing chemo-radiotherapy [11].

In conclusion, DSBA was shown to promote positive hormesis *in vivo*. Mechanistically, Nrf2 activation and downstream GSTP gene induction are confirmed as molecular players of this selenohormetic effect.

## Acknowledgments

BD is a fellow of research supported by the FIRC-AIRC grant program for young investigators.

The study was supported in part by the University of Perugia grant program “Ricerca di base”.

## Author Contributions

**Conceptualization:** Bartolini Desirée, Wang Yanzhong, Giustarini Daniela, Wang Gavin Y.

**Data curation:** Bartolini Desirée, Giustarini Daniela, Wang Gavin Y.

**Formal analysis:** Bartolini Desirée and Wang Gavin Y

**Methodology:** Bartolini Desirée, Wang Yanzhong, Jie Zhang, Torquato Pierangelo and Giustarini Daniela

**Supervision:** Rossi Ranieri, Galli Francesco, Tew Kenneth D. and Townsend, Danyelle M.

**Validation:** Bartolini Desirée and Wang Yanzhong

**Resources:** Galli Francesco, Tew Kenneth D. and Rossi Ranieri

**Writing – original draft:** Bartolini Desirée and Galli Francesco

## References

1. Bartolini D, Sancineto L, Fabro de Bem A, Tew KD, Santi C, et al. (2017) Selenocompounds in Cancer Therapy: An Overview. Adv Cancer Res 136: 259–302.

2. Galli F (2007) Interactions of polyphenolic compounds with drug disposition and metabolism. Curr Drug Metab 8: 830–838.

3. Bartolini D, Piroddi M, Tidei C, Giovagnoli S, Pietrella D, et al. (2015) Reaction kinetics and targeting to cellular glutathione S-transferase of the glutathione peroxidase mimetic PhSeZnCl and its D,Lpolylactide microparticle formulation. Free Radic Biol Med 78: 56–65.

4. Bartolini D, Commodi J, Piroddi M, Incipini L, Sancineto L, et al. (2015) Glutathione S-transferase pi expression regulates the Nrf2-dependent response to hormetic diselenides. Free Radic Biol Med 88: 466–480.

5. Nikawa T, Schuch G, Wagner G, Sies H (1994) Interaction of albumin-bound ebselen with rat liver glutathione S-transferase and microsomal proteins. Biochem Mol Biol Int 32: 291–298.

6. Nikawa T, Schuch G, Wagner G, Sies H (1994) Interaction of ebselen with glutathione S-transferase and papain in vitro. Biochem Pharmacol 47: 1007–1012.

7. Bartolini D, Galli F (2016) The functional interactome of GSTP: A regulatory biomolecular network at the interface with the Nrf2 adaption response to oxidative stress. J Chromatogr B Analyt Technol Biomed Life Sci 1019: 29–44.

8. Wang Y, Liu L, Pazhanisamy SK, Li H, Meng A, et al. (2010) Total body irradiation causes residual bone marrow injury by induction of persistent oxidative stress in murine hematopoietic stem cells. Free Radic Biol Med 48: 348–356.

9. Rossi R, Giustarini D, Colombo G, Milzani A, Dalle-Donne I (2009) Evidence against a role of ketone bodies in the generation of oxidative stress in human erythrocytes by the application of reliable methods for thiol redox form detection. J Chromatogr B Analyt Technol Biomed Life Sci 877: 3467–3474.

10. Wang Y, Liu L, Zhou D (2011) Inhibition of p38 MAPK attenuates ionizing radiation-induced hematopoietic cell senescence and residual bone marrow injury. Radiat Res 176: 743–752.

11. Xiao X, Luo H, Vanek KN LaRue, AC, Schulte BA, et al. (2015) Catalase inhibits ionizing radiation-induced apoptosis in hematopoietic stem and progenitor cells. Stem Cells Dev 24: 1342–1351.

12. Wang Y, Schulte BA, LaRue AC, Ogawa M, Zhou D (2006) Total body irradiation selectively induces murine hematopoietic stem cell senescence. Blood 107: 358–366.

13. He X, Yang A, McDonald DG, Riemer EC, Vanek KN, et al. (2017) MiR-34a modulates ionizing radiation-induced senescence in lung cancer cells. Oncotarget 8: 69797–69807.

14. Zhang J, Shibata A, Ito M, Shuto S, Ito Y, et al. (2011) Synthesis and characterization of a series of highly fluorogenic substrates for glutathione transferases, a general strategy. J Am Chem Soc 133: 14109–14119.

15. Yang A, Qin S, Schulte BA, Ethier SP, Tew KD, et al. (2017) MYC Inhibition Depletes Cancer Stem-like Cells in Triple-Negative Breast Cancer. Cancer Res 77: 6641–6650.

16. Tebay LE, Robertson H, Durant ST, Vitale SR, Penning TM, et al. (2015) Mechanisms of activation of the transcription factor Nrf2 by redox stressors, nutrient cues, and energy status and the pathways through which it attenuates degenerative disease. Free Radic Biol Med 88: 108–146.

17. Olm E, Fernandes AP, Hebert C, Rundlof AK, Larsen EH, et al. (2009) Extracellular thiol-assisted selenium uptake dependent on the x (c)-cystine transporter explains the cancer-specific cytotoxicity of selenite. Proc Natl Acad Sci U S A 106: 11400–11405.

18. De Nicola M, Ghibelli L (2014) Glutathione depletion in survival and apoptotic pathways. Front Pharmacol 5: 267.

19. Tew KD (1994) Glutathione-associated enzymes in anticancer drug resistance. Cancer Res 54: 4313–4320.

20. Zhang J, Ye ZW, Gao P, Reyes L, Jones EE, et al. (2014) Glutathione S-transferase P influences redox and migration pathways in bone marrow. PLoS One 9: e107478.

21. Markovic J, Garcia-Gimenez JL, Gimeno A, Vina J, Pallardo FV (2010) Role of glutathione in cell nucleus. Free Radic Res 44: 721–733.

22. Li H, Wang Y, Pazhanisamy SK, Shao L, Batinic-Haberle I, et al. (2011) Mn (III) meso-tetrakis-(Nethylpyridinium-2-yl) porphyrin mitigates total body irradiation-induced long-term bone marrow suppression. Free Radic Biol Med 51: 30–37.

